# Population genomics of two invasive mosquitoes (*Aedes aegypti* and *Aedes albopictus*) from the Indo-Pacific

**DOI:** 10.1101/2020.03.15.993055

**Authors:** Thomas L Schmidt, Jessica Chung, Ann-Christin Honnen, Andrew R Weeks, Ary A Hoffmann

## Abstract

The arbovirus vectors *Aedes aegypti* (yellow fever mosquito) and *Ae. albopictus* (Asian tiger mosquito) are both common throughout the Indo-Pacific region, where 70% of global dengue transmission occurs. For *Ae. aegypti* all Indo-Pacific populations are invasive, having spread from an initial native range of Africa, while for *Ae. albopictus* the Indo-Pacific includes invasive populations and those from the native range: putatively, India to Japan to Southeast Asia. This study analyses the population genomics of 480 of these mosquitoes sampled from 27 locations in the Indo-Pacific. We investigated patterns of genome-wide genetic differentiation to compare pathways of invasion and ongoing gene flow in both species, and to compare invasive and native-range populations of *Ae. albopictus*. We also tested landscape genomic hypotheses that genetic differentiation would increase with geographical distance and be lower between locations with high connectivity to human transportation routes, the primary means of dispersal at these scales. We found that genetic distances were generally higher in *Ae. aegypti*, with Pacific populations the most highly differentiated. The most differentiated *Ae. albopictus* populations were in Vanuatu, Indonesia and Sri Lanka, the latter two representing potential native-range populations and potential cryptic subspeciation respectively. Genetic distances in *Ae. aegypti* increased with geographical distance, while in *Ae. albopictus* they decreased with higher connectivity to human transportation routes. Contrary to the situation in *Ae. aegypti*, we found evidence of long-distance *Ae. albopictus* colonisation events, including colonisation of Mauritius from East Asia and of Fiji from Southeast Asia. These direct genomic comparisons indicate likely differences in dispersal ecology in these species, despite their broadly sympatric distributions and similar use of human transport to disperse. Our findings will assist biosecurity operations to trace the source of invasive material and for biocontrol operations that benefit from matching genetic backgrounds of released and local populations.

**Author Summary:** The mosquitoes *Ae. aegypti* and *Ae. albopictus* are highly invasive and transmit dengue and other arboviruses. This study investigates the genetics of these mosquitoes in the Indo-Pacific region, where 70% of global dengue transmission occurs and where both species have established widespread invasions by hitch-hiking on human transport vessels. We compared patterns of genetic differentiation to determine the pathways these species have taken while spreading through the Indo-Pacific, and to better understand how they disperse. We sequenced DNA from 480 mosquitoes sampled from 27 locations in the Indo-Pacific, and found many genetic differences between the two species. Populations of *Ae. aegypti*, which is not native to the region, tended to be genetically different from each other, and populations in the Pacific Ocean were particularly divergent. *Aedes albopictus* populations were generally more similar to each other, though genetically different populations in Sri Lanka and Indonesia point to these having a different history to other populations. Genetic differences between *Ae. aegypti* populations were larger when populations were geographically distant, while differences between *Ae. albopictus* populations were larger when populations likely had limited access to human transportation. These results will help improve strategies for controlling these species and stopping their spread around the world.

## Introduction

The Indo-Pacific region, here defined as encompassing the Indian Ocean, the western and central Pacific Ocean, and the coastal territories therein, is the site of 70% of global dengue transmission [1]. Infections are vectored by two mosquito species, *Aedes aegypti* (yellow fever mosquito) and *Ae. albopictus* (Asian tiger mosquito) [2,3], both of which have established widespread invasions in the region. *Aedes aegypti*, the primary regional vector, has a putative native range of West Africa [4] and is thought to have invaded the Indian Ocean via the Mediterranean [5], while its invasion history in the western Pacific remains unclear but likely happened in the ∼18-19^th^ c. [6]. *Aedes albopictus* has a hypothesised native range that includes Japan, China, northern India, and parts of Southeast Asia [7]. The southern extent of this range remains unclear and may extend as far as Indonesia and New Guinea [7–9]. Regional expansion of *Ae. albopictus* is thought to have occurred more recently than *Ae. aegypti*, with colonisation likely involving successive waves [8,10].

Like many invasive taxa, *Ae. aegypti* and *Ae. albopictus* (hereafter *Aedes* spp.) exhibit stratified dispersal. This consists of two distinct processes: short-range, active dispersal and long-range, passive dispersal [11]. Active dispersal of *Aedes aegypti* is generally highly localised, as measured by both traditional mark-release-recapture methods (8 – 199 m/gen [12]) or current methods using genomic estimates of relatedness (33 – 131 m/gen [13]). Similar observations have been made of *Ae. albopictus* using mark-release-recapture [14]. At the metropolitan scale (< 50 km), passive dispersal in *Aedes* spp. has been observed directly [15] or inferred from the spatial distribution of close kin [16,17]. Dispersal at broader scales such as between cities is almost exclusively by passive transportation on ships, aircraft and land vehicles [18–21].

The unrestricted spread of *Aedes* spp. has many adverse consequences. Both species are presently continuing to colonise new tropical, subtropical and, for *Ae. albopictus*, temperate regions [22], and future expansions are expected in the face of increased urbanisation and climate change [23]. In addition to range expansions, dispersal between *Aedes* spp. populations can spread advantageous alleles that reduce the efficacy of local control programs [24]. Incursions of these species are frequently linked to vessels such as ships and planes [19,20,25], and when source locations can be identified they are often far from the point of collection [20,25].

Understanding the forces that influence dispersal, gene flow, and population structure in *Aedes* spp. can help efforts to prevent, control, and prepare for future threats from these species. As well as being a critical focus for dengue control efforts, the Indo-Pacific presents a useful location to investigate these processes due to the broad sympatry of *Aedes* spp. within the region. Investigating the genetics of sympatric taxa can help link patterns of genetic differentiation to the forces that shape population structure [26,27]. For instance, when taxa share a common area but have contrasting patterns of differentiation, this may indicate intrinsic differences in how they disperse [28–30], while similar dispersal behaviour can produce similar patterns of differentiation [31].

Here we investigate these processes in *Aedes* spp. from the Indo-Pacific region at spatial scales from ∼20 km (inter-city) to ∼16,000 km (trans-oceanic). Dispersal at these scales largely proceeds along the network of shipping routes that link each inhabited coastal location with every other location (Fig 1). As *Aedes* spp. are thought to make use of the same vessel types for transportation, their dispersal may take place along similar pathways, which may produce similar patterns of genetic structure [31,32]. However, there are several reasons why different patterns might be observed. *Aedes albopictus* may be able to survive transportation over longer distances than *Ae. aegypti* by undergoing quiescence and diapause [33–35] which confers resistance to cold and desiccation [36]. *Aedes aegypti* does not undergo diapause but can make use of egg quiescence to survive adverse conditions [35]. Heterogeneous histories of colonisation may also produce differences in structure. At broad scales, this relates to differences between the two species in the timing and direction of invasions. At finer scales, stochastic processes during colonisation can produce rapid change in allele frequencies [37]. For any of the above processes, genetic structure can be resistant to ‘erosion’ via gene flow if local population densities are high [38] as they generally are for *Aedes* spp.

**Fig 1.**
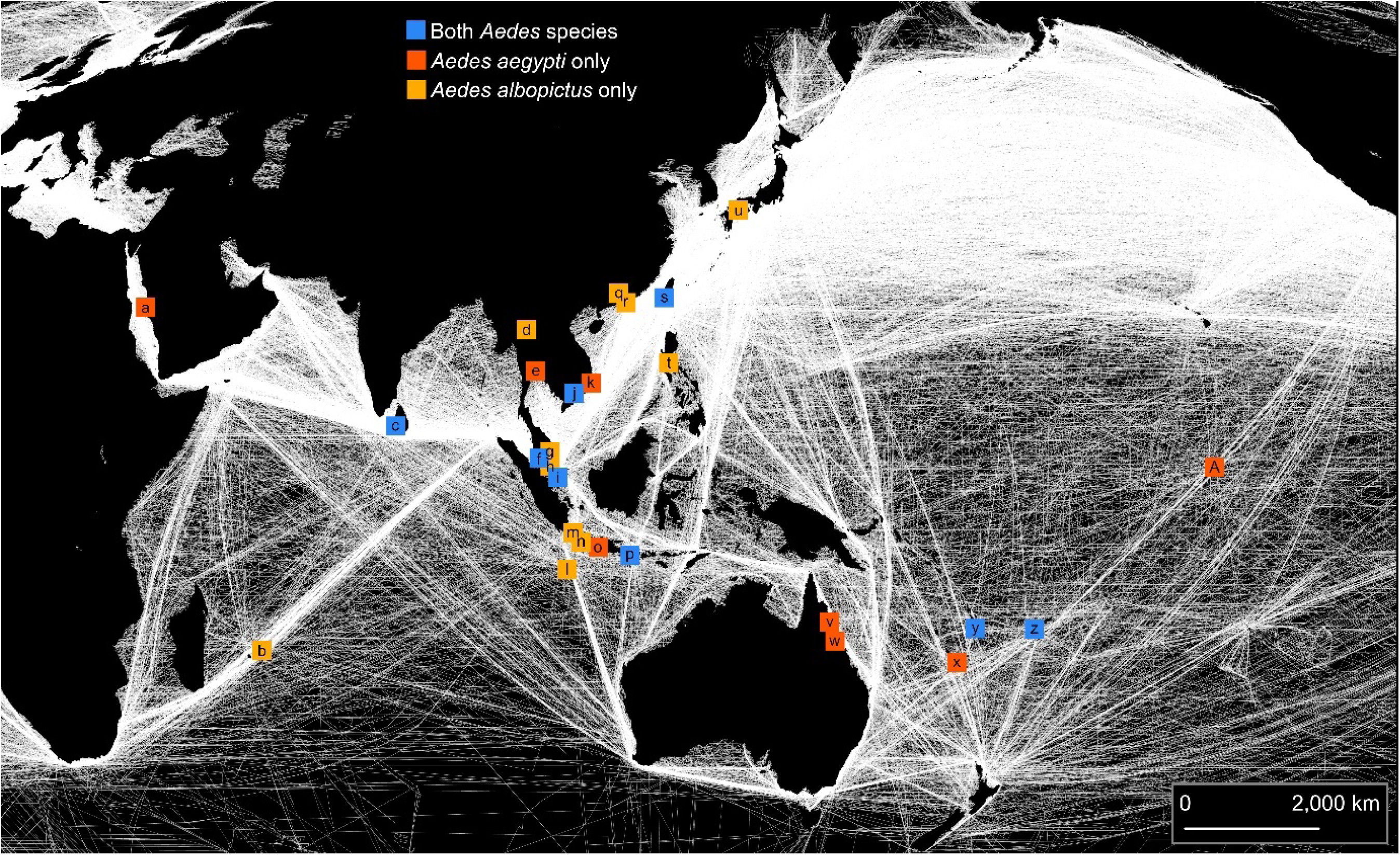
Sampling locations of *Aedes aegypti* and *Aedes albopictus*. White lines indicate density of shipping routes. The map uses a Mollweide projection with a central meridian of 120°E. The map was produced in ARCMAP 10.5.1 using shipping route data made available from Halpern et al. [39]. Sampling locations are indicated by letters and are described in Tables 1 and 2.

This study uses high-resolution markers to compare patterns of broad-scale genetic structure in *Aedes* species. Few such comparative studies have been conducted [40,41], and none have used high-resolution molecular markers, which have greater demonstrated power to distinguish between *Aedes* populations [42]. We also test landscape genomic hypotheses of isolation by distance [43] and gene flow via connectivity to human transportation routes. These analyses provide a means of disentangling the putative influences of human transportation routes which may direct gene flow, high local population density following invasion which may limit the effects of gene flow, and diapause in *Ae. albopictus* which may permit gene flow between more distant populations than for *Ae. aegypti*. Recently, single-species studies using high-resolution molecular markers have reported strong divergence among *Ae. aegypti* populations [5,42] and weaker divergence among *Ae. albopictus* populations [44,45], though highly-differentiated *Ae. albopictus* populations have been recorded [46]. As broad-scale studies have typically included a greater number of samples from European, African and American *Aedes* spp. populations than from Indo-Pacific populations [5,44,45,47], this study also fills an important geographical gap in the population genomics literature on *Aedes* spp..

**Table 1.** Details of *Aedes aegypti* sampled from 16 populations. See Fig 1 for map id locations. See Materials and Methods for details regarding filtering and calculation of connectivity indices.

| map id<br>(see Fig 1) | sample | location(s) | country | year(s)<br>collected | no. genotypes<br>after filtering | connectivity<br>(marine) | connectivity<br>(aerial) |
| --- | --- | --- | --- | --- | --- | --- | --- |
| a | Jeddah | Various locations in Jeddah | Saudi Arabia | 2018 | 18 | 7.92 | 194 |
| c | Colombo | Kiribathgoda | Sri Lanka | 2017 | 12 | 9.62 | 101 |
| e | Bangkok | Seven locations in Bangkok | Thailand | 2016 | 18 | 0.17 | 326 |
| f | Kuala Lumpur | Bandar Sunway/UKM Specialist Centre | Malaysia | 2015/2017 | 18 | 3 | 257 |
| i | Singapore | Various locations in Singapore | Singapore | 2015 | 15 | 2.64 | 408 |
| j | Ho Chi Minh City | Various locations in Ho Chi Minh City | Vietnam | 2014 | 18 | 0.43 | 109 |
| k | Nha Trang | Hon Mieu Island | Vietnam | 2016 | 11 | 5.26 | 7 |
| o | Yogyakarta | Various locations in Yogyakarta, Java | Indonesia | 2014 | 13 | 0.44 | 29 |
| p | Bali | Near Denpasar Airport, Bali | Indonesia | 2016/2017 | 18 | 1.24 | 100 |
| s | Taiwan | Various locations in Southwest Taiwan | Republic of China | 2016 | 8 | 7.25 | 65 |
| v | Cairns | Six locations in urban Cairns | Australia | 2015 | 18 | 1.86 | 56 |
| w | Townsville | Near Townsville Airport | Australia | 2014 | 10 | 2.25 | 16 |
| x | New Caledonia | Various locations in Nouméa | New Caledonia | 2018 | 13 | 1.24 | 13 |
| y | Vanuatu | Various locations in Efate | Vanuatu | 2018 | 14 | 0.71 | 28 |
| z | Fiji | Various locations in Nadi | Fiji | 2018 | 11 | 0.85 | 43 |
| A | Kiribati | Various locations in Kiribati | Kiribati | 2018 | 9 | 0.8 | 3 |

**Table 2.** Details of *Aedes albopictus* sampled from 20 populations. See Fig 1 for map id locations. See Materials and Methods for details regarding filtering and calculation of connectivity indices. Colombo-1 was treated as a distinct population due to divergences (see text), and omitted (*) from landscape genomic analyses.

| map id<br>(see Fig 1) | sample name | location(s) | country | year(s)<br>collected | no. genotypes<br>after filtering | connectivity<br>(marine) | connectivity<br>(aerial) |
| --- | --- | --- | --- | --- | --- | --- | --- |
| b | Mauritius | Near Chamarel village | Mauritius | 2017 | 7 | 2.68 | 41 |
| c | Colombo-1 | Delgoda | Sri Lanka | 2017 | 7 | * | * |
| c | Colombo-2 | Delgoda and Kiribathgoda | Sri Lanka | 2017 | 8 | 9.62 | 101 |
| d | Chiang Mai | Near Wachirathan Waterfall | Thailand | 2017 | 5 | 0 | 48 |
| f | Selangor | Kuala Lumpur and Gombak, Selangor | Malaysia | 2016 | 11 | 3 | 257 |
| g | Pahang | Kuantan | Malaysia | 2015 | 18 | 3.2 | 5 |
| h | Johor | Johor Baru | Malaysia | 2016 | 18 | 3.28 | 18 |
| i | Singapore | Near Bukit Timah Nature Reserve | Singapore | 2017 | 14 | 2.07 | 408 |
| j | Ho Chi Minh City | Various locations in Ho Chi Minh City | Vietnam | 2017 | 18 | 0.44 | 109 |
| l | Christmas Island | Near Christmas Island International Airport | Australia | 2017 | 7 | 1.91 | 2 |
| m | Jakarta | Various locations in Jakarta, Java | Indonesia | 2016 | 17 | 1.09 | 185 |
| n | Bandung | Various locations in Bandung, Java | Indonesia | 2017 | 14 | 0.53 | 21 |
| p | Bali | Various locations in Bali | Indonesia | 2016/2017 | 17 | 1.24 | 100 |
| q | Guangzhou | 13 locations in Guangzhou | China | 2015 | 16 | 0.24 | 338 |
| r | Hong Kong | Various locations in Hong Kong | China | 2016 | 16 | 3.87 | 355 |
| s | Taiwan | Various locations in Southwest Taiwan | Republic of China | 2016 | 15 | 7.24 | 65 |
| t | Manila | Manila, Quezon City and Makati City | Philippines | 2017 | 6 | 1.56 | 193 |
| u | Matsuyama | Near Ehime University | Japan | 2017 | 11 | 1.97 | 12 |
| y | Vanuatu | Various locations in Efate | Vanuatu | 2018 | 15 | 0.71 | 28 |
| z | Fiji | Various locations in and near to Nadi | Fiji | 2018 | 16 | 0.84 | 43 |

We found clear differences between the spatial genetic structure of *Ae. aegypti* and *Ae. albopictus*, and our landscape genomic analyses indicated that *Ae. aegypti* populations were generally structured by space, and *Ae. albopictus* populations by connectivity to human transport routes. These results may reflect important differences in the capacity of *Aedes* spp. to disperse and to invade new regions, and may also reflect the limited power of gene flow to erode existing structure in species with high census sizes. The findings of this study will assist biosecurity operations that aim to trace the source of invasive material [20,48] and for biocontrol operations that benefit from matching genetic backgrounds of released and local populations [49].

## Materials and Methods

### Sample collection, genotyping, processing and subsampling

*Aedes aegypti* were sampled from 16 locations in the Indo-Pacific region, across a ∼16,000 km range from Jeddah, Saudi Arabia to Kiribati (Fig 1, Table 1). *Aedes albopictus* were sampled from 19 locations ranging from Mauritius to Fiji to Matsuyama, Japan (Fig 1, Table 2). We considered mosquitoes collected within the same city to be from the same population, as structuring within these scales is typically weak (e.g. [17]). The smallest distance between two distinct populations was ∼20 km, between Singapore and Johor, Malaysia. Among populations within the same country, the smallest distance was ∼130 km, between Guangzhou and Hong Kong in China.

Mosquitoes were genotyped using a pipeline that has been described elsewhere [20] and is described here in Appendix S1. Briefly, we used the double-digest restriction site-associated (ddRAD) sequencing protocol for *Ae. aegypti* [42] to obtain sequence reads, which were processed in Stacks v2.0 [50]. We used BOWTIE V2.0 [51] to align reads to the *Ae. aegypti* [52] and *Ae. Albopictus* [53] mitochondrial genome (mtDNA) assemblies. Reads that did not align to the mtDNA assemblies were aligned to the nuclear assemblies AaegL5 [54] and AaloF1 [55] to obtain nuclear genotypes for *Ae. aegypti* and *Ae. albopictus* respectively. This ensured that the nuclear genotypes would not be confounded by transpositions of mitochondrial DNA into the nuclear genome (NUMTs), a common occurrence in *Aedes* spp. [56]. As some population samples contained more close kin than others, we removed individuals in order of missing data so that no first order kin pairs remained and no population had greater than 18 genotypes, following the procedure described in Appendix S1.

We retained 224 *Ae. aegypti* and 256 *Ae. albopictus* genotypes. We used the Stacks program REF_MAP to build catalogs containing all mosquito genotypes from each species, from which we called single nucleotide polymorphism (SNP) genotypes at RAD stacks using default Stacks settings. Only SNPs scored in at least 75% of genotypes from each population were used in analyses.

### Individual-level genetic structure

We investigated genetic structure among individual mosquito genotypes using three methods: discriminant analysis of principal components (DAPC [57]); sparse non-negative matrix factorisation (sNMF [58]); and PCA-UMAP [59], which combines principal components analysis (PCA) with uniform manifold approximation and projection (UMAP [60]). For these analyses, we filtered the 224 *Ae. aegypti* and 256 *Ae. albopictus* genotypes to retain biallelic SNPs with < 20% missing data, with minimum depth of coverage of 3, and maximum depth of coverage of 45. Genotypes were phased and missing data imputed in BEAGLE V4.1 [61] using default settings and 10 iterations.

DAPC was run in the R package *adegenet* [62]. In DAPC, each genotype is assigned to one of *K* genetic clusters. We ran the ‘find.clusters’ algorithm to determine the optimal *K* for 1 ≤ *K* ≤ N, with N equal to the number of populations (16 for *Ae. aegypti*, 20 for *Ae. albopictus*). We ran 1×10^9^ iterations and selected the value of *K* with the lowest Bayesian information criterion (BIC). We then repeated the above analysis, selecting the value of *K* with the lowest Akaike information criterion (AIC), a less conservative measure than BIC.

We ran sNMF in the R package LEA [63]. This analysis estimates individual ancestry coefficients, assuming that individual genotypes are produced from the admixture of *K* ancestral lineages, where *K* is unknown *a priori*. To determine which value of *K* was optimal for summarising the variation in each species, we set 1 ≤ *K* ≤ N and ran 100 iterations of the sNMF algorithm for each *K*, selecting the *K* with the lowest cross-entropy across all runs. For the chosen *K* we selected the iteration with the lowest cross-entropy for visual presentation.

To generate PCA-UMAPs, we adapted code provided in Diaz-Papkovich et al. [59]. UMAP provides a means of projecting high-dimensional data onto two-dimensional space, and has advantages over other dimensionality reduction techniques in that it better preserves the global data structure between clusters in reduced dimensions. Combining UMAP and PCA has been shown to produce optimum definition of population clusters [59]. PCA-UMAPs used the first 5 principal components of the PCA, which were projected in two dimensions via UMAP using 50 neighbours and a minimum distance of 0.5.

### Population-level genetic structure

For investigations of population-level genetic structure, we were cautious to process and filter the data so that uneven population sample size (*n*) would not bias analyses, while still including as many genotypes as possible. To balance these aims, we subsampled each dataset ten times, sampling (with replacement) from each population a number of genotypes (five) equal to the minimum *n* among all populations. For each subsample we used the Stacks program POPULATIONS to calculate F_ST_’ [64] between each pair of populations, and to calculate mean observed heterozygosity (H_O_), nucleotide diversity (π), number of private alleles and proportion of missing data for each population. We used the results from the ten subsamples to calculate the mean and 95% confidence intervals of each estimated parameter.

For genome-wide SNP datasets, genetic differentiation can be estimated with five genotypes [65]. To determine whether using only five genotypes in each subsample affected our F_ST_’ estimates, we took ten subsamples of 16 individuals from the *Ae. aegypti* populations in Bali, Bangkok, Jeddah and Kuala Lumpur, and calculated F_ST_’ means and 95% confidence intervals as above. We compared these results with those calculated with five genotypes.

Analysis of the data indicated a deep genetic division within the Sri Lankan *Ae. albopictus* sample (see Fig 2ii and Results: Individual-level genetic structure). Seven of the 15 genotypes formed a group highly divergent from all others, including the other Sri Lankan genotypes sampled from the same region of Colombo. This did not appear to be an artefact of sampling close kin; only a single half-sib dyad was present in the 7 genotypes, a similar incidence as in other populations. We therefore treated the Sri Lankan *Ae. albopictus* sample as two separate populations, denoted as Colombo-1 (highly divergent) and Colombo-2. We investigated this divergence further by calculating the folded allele frequency spectrum for each *Ae. Albopictus* population in VCFTOOLS [66], using the ten subsamples of five genotypes to calculate means and 95% confidence intervals.

**Fig 2.**
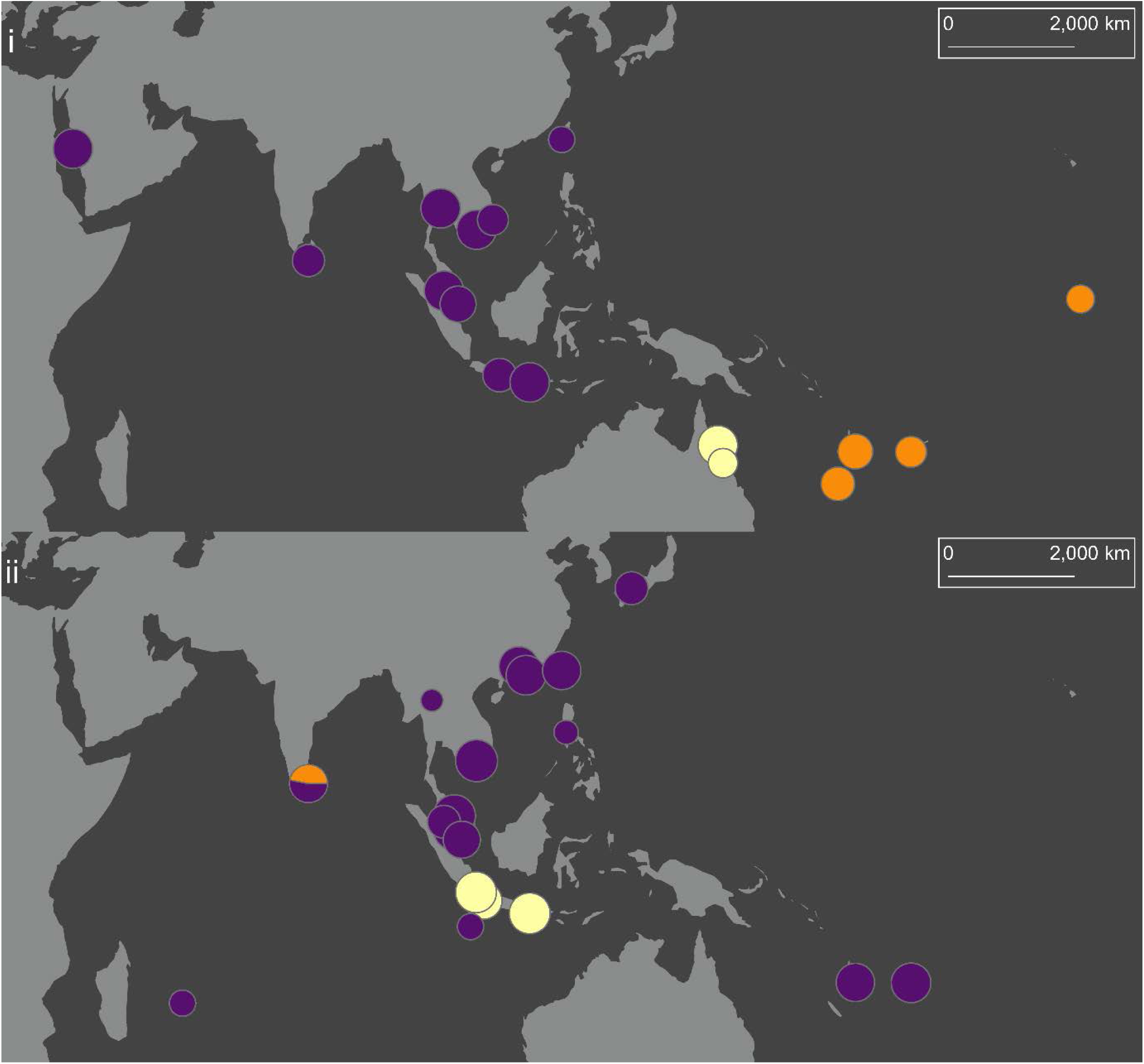
DAPCs of *Aedes aegypti* (i) and *Aedes albopictus* (ii) using lowest BIC (*K* = 3). Colours indicate cluster membership. Circles are sized relative to population sample size. The map uses a Mollweide projection with a central meridian of 120°E. The map was produced in ARCMAP 10.5.1, using shapefiles made available by www.naturalearthdata.com.

### Landscape genomics

We investigated the geographical structuring of genetic variation in *Aedes* spp. using distance-based redundancy analysis (dbRDA), performed in the R package *vegan* [67]. These analyses tested two hypotheses: that genetic differentiation would increase with geographical distance (i.e. isolation by distance), and that genetic differentiation would decrease with greater connectivity to shipping or aerial transportation routes. We omitted the outlier *Ae. albopictus* subpopulation Colombo-1 from analyses (see Results: Individual-level genetic structure). Mean F_ST_’ estimates among the 16 *Ae. aegypti* and 19 *Ae. albopictus* populations were used as a measure of genetic differentiation.

Our measure for connectivity to shipping routes was derived from previously published raster data of 799,853 commercial shipping tracks from October 2004 to October 2005 (Halpern et al., 2008; see Supporting Online Material therein for full description). We used ARCMAP 10.5.1 [68] to calculate the average density of shipping tracks within a 200 km radius from each population sampling point, so that the inland site at Chiang Mai scored 0 while Jeddah, Colombo-2, and Taiwan received the highest scores (see Fig 1). For connectivity to aerial transportation routes, we used data from the OpenFlights Routes Database (https://openflights.org/data.html#route), which lists all airline routes active at June 2014. For each sampled population, we used the total number of active flight routes at the largest international airport of that city or town. This gave values of between 2 (Christmas Island) to 408 (Singapore). Shipping and aerial connectivity scores for each population are listed in Tables 1 and 2. No correlation between shipping and aerial connectivity scores were observed for either *Ae. aegypti* (R^2^ = 0.003, P = 0.838) or *Ae. albopictus* (R^2^ < 0.001, P = 0.938) populations.

The connectivity of each population pair to the shipping and aerial networks was calculated as the average of their connectivity scores for that network type. These average scores were used to construct pairwise connectivity matrices of each transportation type for each species. As independent variables in a dbRDA must be rectangular, these matrices were transformed into principle coordinates using the function *pcnm*. We likewise calculated geographical distance matrices for each species, and transformed these into principle coordinates.

To test for isolation by distance in each species, we built dbRDAs using the scaled vectors of geographical distance as the independent variable and F_ST_’ as the dependent variable. To test for effects of connectivity to transportation networks, we first determined scaled geographical distance vectors with which to condition the model, by building dbRDAs where each vector was treated as a separate independent variable, and selecting vectors with P < 0.01. We assessed shipping and aerial connectivity separately. We built dbRDAs using the scaled vectors of connectivity as the independent variable and F_ST_’ as the dependent variable, with the model conditioned using the significant vectors for geographical distance. All dbRDAs were built using the function *capscale*. Marginal significance of variables was assessed using *anova.cca* with 9,999 permutations.

Once transformed into principal coordinates, geographical and connectivity variables ceased to indicate directionality, and thus dbRDAs could only ascertain significance of associations and not the direction of these associations. To determine directionality, we ran linear regressions on the normalised pairwise scores of these variables against normalised F_ST_’ estimates, and used the regression coefficients as indicators of directionality. These analyses were not used to determine significance of variables due to non-independence among pairwise terms.

## Results

### Individual-level genetic structure

We retained 54,296 SNPs for analysis of the 224 *Ae. aegypti* (Table 1) and 40,016 SNPs for analysis of the 256 *Ae. albopictus* (Table 2), with mean read depths of coverage of 17.67 (s.d. 5.31) and 25.23 (s.d. 7.49) respectively. The three analyses of genetic structure among individual genotypes, DAPC, sNMF, and PCA-UMAP, were consistent in their findings, and indicated several broad differences in genetic structure among *Aedes* spp. in the Indo-Pacific.

DAPC determined that *K* = 3 was the optimal number of clusters for both *Ae. aegypti* and *Ae. albopictus* when lowest BIC (Fig 2) was used for cluster detection. For *Ae. aegypti*, DAPC partitioned genotypes into three spatial groups representing the Pacific Islands (orange), Australia (cream), and all central and Western populations (purple) (Fig 2i). DAPC of *Ae. albopictus* (Fig 2ii) showed a very different pattern, with a single cluster (purple) present across the entire sampling range. One other cluster (cream) was found in Indonesia, indicating deep differences between these populations and nearby populations like Christmas Island, despite separation distances of only ∼450 km. The final *Ae. albopictus* cluster was found within the Colombo population, consisting of a subset of seven genotypes that were deeply divergent from the other eight Colombo genotypes. In all further analyses the Colombo genotypes were treated as two populations, denoted as Colombo-1 (highly divergent) and Colombo-2. As BIC is a conservative measure, these results represent broad genetic groupings, with more detailed variation described in sNMFs (Fig 3), PCA-UMAPs (Figs 4 and 5), and DAPC using lowest AIC (S1 Fig A).

**Fig 3.**
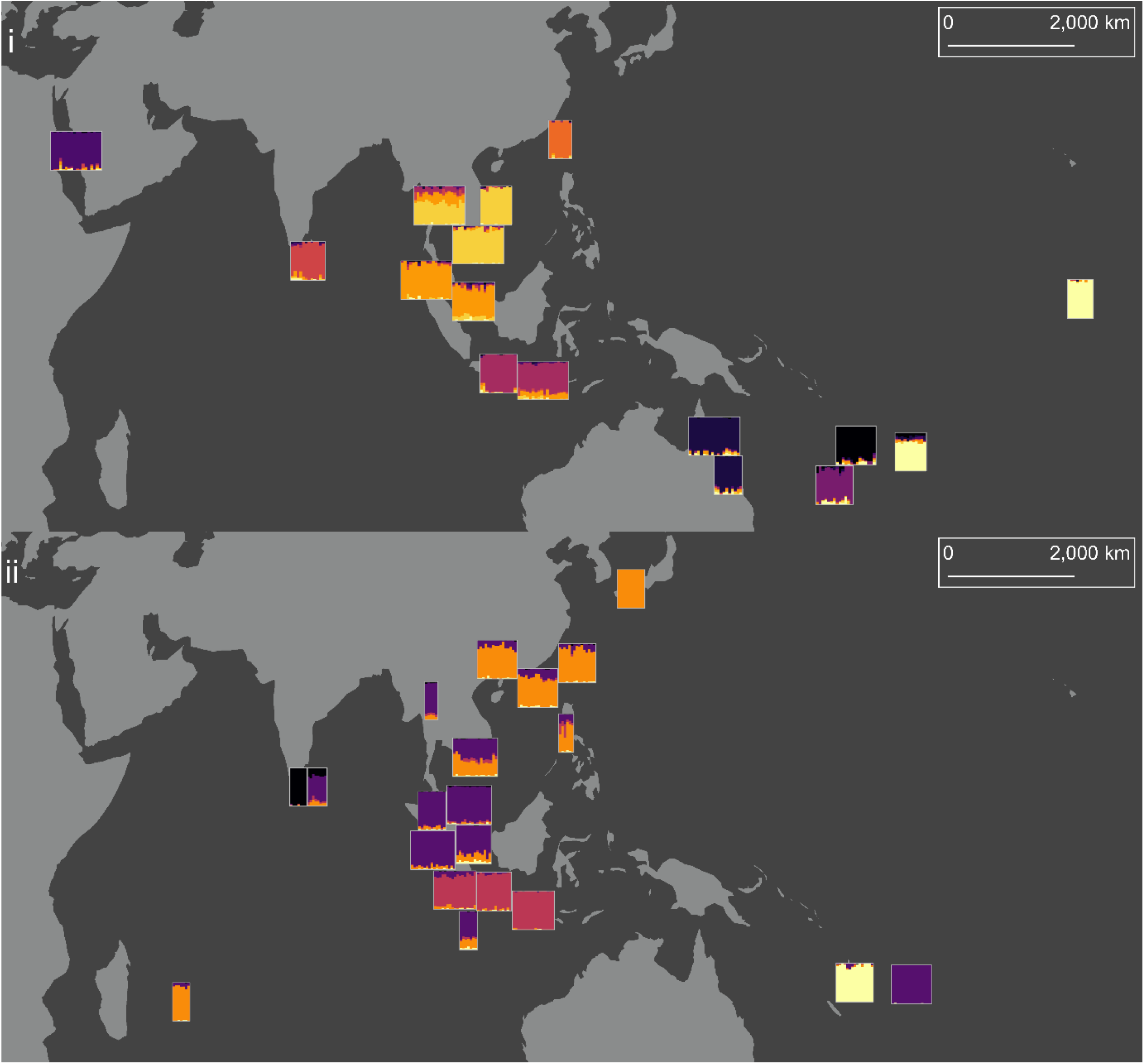
sNMFs of *Aedes aegypti* (i) and *Aedes albopictus* (ii). Number of ancestral lineages (*K*) was 10 for *Ae. aegypti* and 5 for *Ae. albopictus*. Each population is indicated as a rectangle, with each genotype a vertical line made of between 1 and *K* colours. Colours indicate ancestral lineages. The map uses a Mollweide projection with a central meridian of 120°E. The map was produced in ARCMAP 10.5.1, using shapefiles made available by www.naturalearthdata.com.

**Fig 4.**
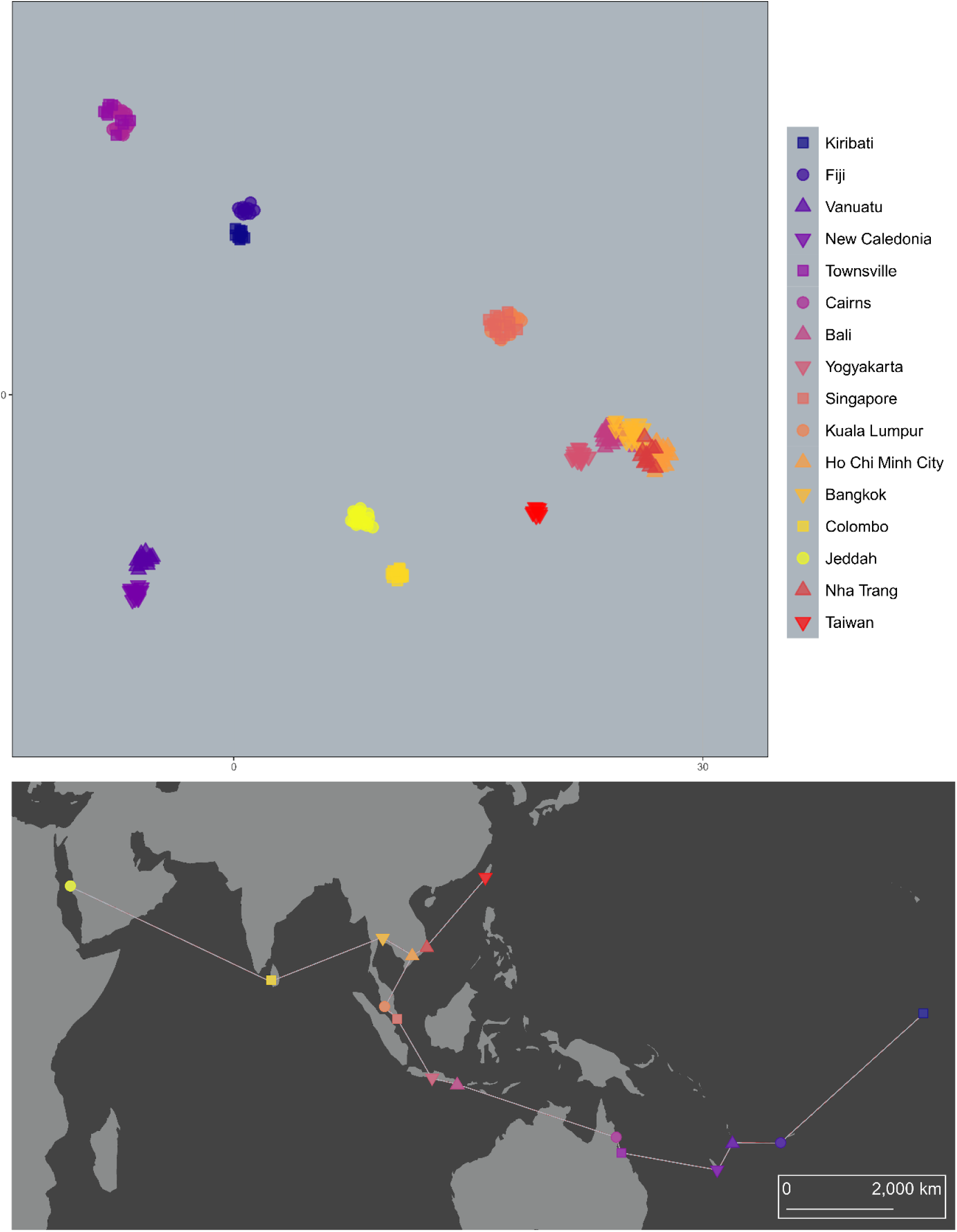
PCA-UMAP of 224 *Aedes aegypti* genotypes. Genotypes are coloured by population, with similar colours given to geographically proximate populations.

**Fig 5.**
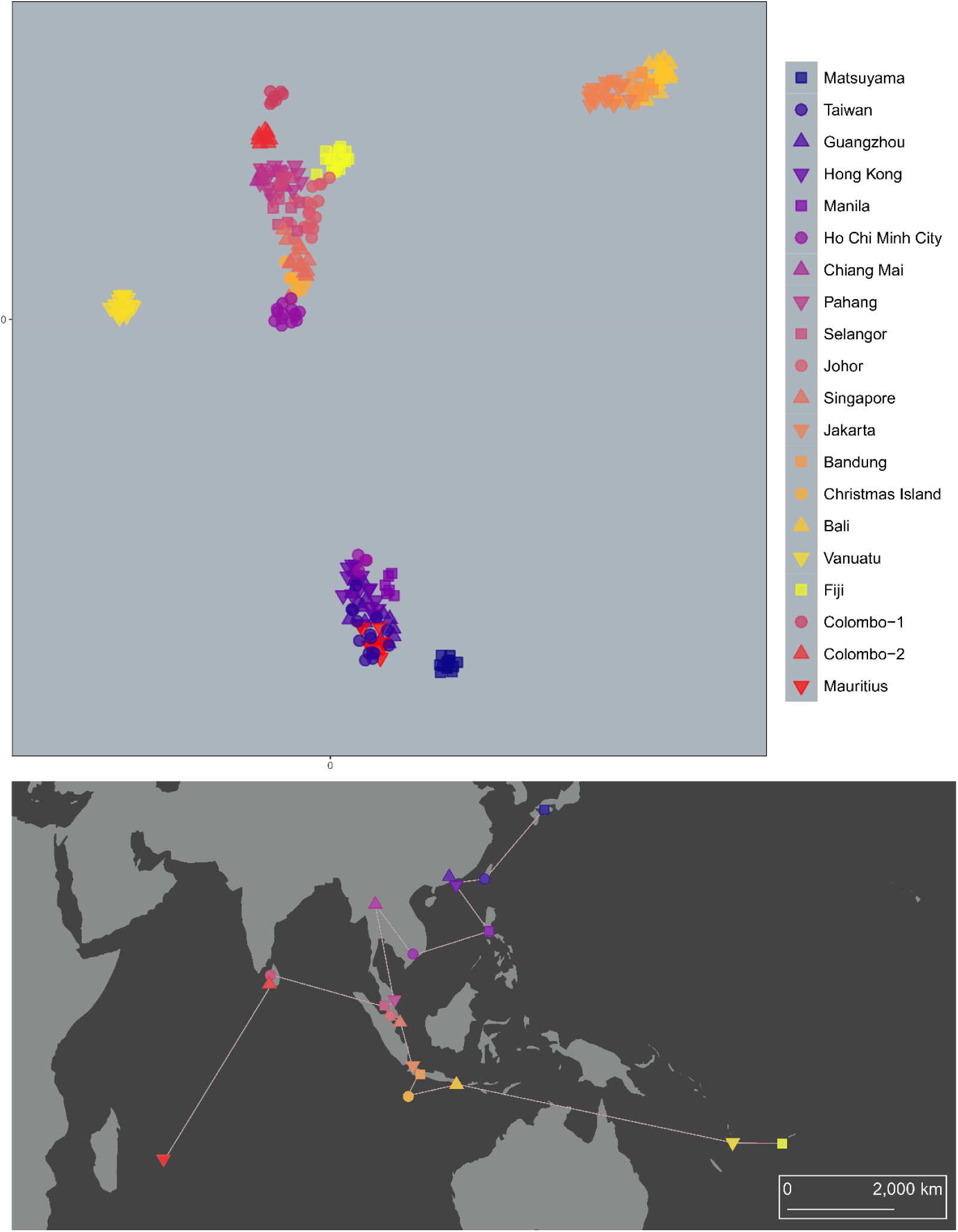
PCA-UMAP of 256 *Aedes albopictus* genotypes. Genotypes are coloured by population, with similar colours given to geographically proximate populations.

The sNMF of *Ae. aegypti* (Fig 3i) found *K* = 10 was an optimal number of ancestral lineages for summarising genetic variation among genotypes. This high number relative to the number of populations (16) largely reflected the high level of structuring between populations, with five of the ten ancestral lineages almost wholly confined to single locations: Jeddah, Colombo, Taiwan, New Caledonia and Vanuatu. Proximate population pairs such as Cairns and Townsville, Kuala Lumpur and Singapore, and Ho Chi Minh City and Nha Trang each had common ancestral lineages, as did Fiji and Kiribati, despite their distance of separation. The most putatively admixed population was Bangkok, which showed genetic similarities with the other Southeast Asian populations, but not with either Taiwan or Colombo.

These results contrasted sharply with those of *Ae. albopictus* (Fig 3ii), for which *K* = 5 was optimal. Fewer ancestral lineages indicated greater genetic similarity among *Ae. albopictus* populations. The interpretation of the spatial distribution of ancestral lineages among *Ae. albopictus* requires reference to the native and invasive ranges of this species. The northernmost native range lineage, in orange, indicated a common heritage among the East Asian populations. This lineage was also dominant in Mauritius. A second native range lineage, in purple, indicated common heritage among Southeast Asian populations from Chiang Mai to Singapore, and was also found in Christmas Island and Fiji, but not Vanuatu. The population at Ho Chi Minh City was made up of roughly even contributions from the two lineages. The Indonesian *Ae. albopictus* genotypes (Fig 3ii, red) of Jakarta, Bandung and Bali were genetically distinct from the East and Southeast Asian native range genotypes.

Treating Colombo *Ae. albopictus* as two separate populations revealed that genotypes from the highly-divergent Colombo-1 population were almost entirely of one ancestral lineage (Fig 3ii, black) found only in Colombo. Colombo-2 was a composite of lineages, including the unique lineage from Colombo and the purple native range lineage.

PCA-UMAP of *Ae. aegypti* (Fig 4) showed genotypes grouping clearly by spatial location, echoing the sNMF results but providing more detailed information on differentiation between similar populations. Genotypes clustered first by population, then by region, with Pacific and Australian genotypes the most distinct. PCA-UMAP indicated similarities between genotypes from Fiji and Kiribati, Vanuatu and New Caledonia, and Singapore and Kuala Lumpur. The westernmost samples from Jeddah and Colombo also showed evidence of similarity. Despite Taiwan’s proximity to Southeast Asia, Taiwanese *Ae. aegypti* were clearly distinct from all others.

PCA-UMAP of *Ae. albopictus* (Fig 5) likewise gave results similar to the sNMF but with more detail. The Matsuyama population was revealed as distinct from others in East Asia, and there was evidence that the Fijian genotypes were most closely related to Johor, Malaysia, and those of Christmas Island were most closely related to Singapore. The Ho Chi Minh City genotypes were split into two clusters, four grouping with the East Asian genotypes and 14 with the Southeast Asian genotypes. This likely reflects Ho Chi Minh City as having ancestry from both of these groups (Fig 3ii). A corresponding split was observed in the DAPC results using lowest AIC (S1 Fig A).

### Population-level genetic structure

Numbers of nuclear SNPs retained for population-level analyses had ranges of 46,151 – 54,647 (*Ae. aegypti*) and 35,984 – 57,639 (*Ae. albopictus*). Pairwise F_ST_’ estimates for *Ae. aegypti* and *Ae. albopictus* are listed in S1 Tables A and B respectively. Mean estimates are given with 95% confidence ranges, which show that pairwise F_ST_’ estimates were generally consistent across the ten subsamples. F_ST_’ confidence intervals in *Ae. aegypti* were on average only 5.3% the size of the mean; for *Ae. albopictus*, they were only 6.4% the size of the mean. Mean H_O_ and π estimates were also mostly consistent among subsamples within populations, and are listed in S1 Tables C & D along with the number of private alleles and proportion of missing data among genotypes in each population.

Fig 6 shows all population pairs with mean pairwise F_ST_’ < 0.03 (a value selected to help visualise patterns), indicating high genetic similarity. Australian *Ae. aegypti* populations were genetically very similar, and there was a network of similarity among Southeast Asian populations (Fig 6i). Indonesian *Ae. albopictus* populations were genetically similar, as were a network of populations covering parts of the native range in East and Southeast Asia as well as Mauritius (Fig 6ii). Taiwan *Ae. aegypti* had the highest mean F_ST_’ from all other populations (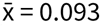; s.d. = 0.016). Colombo-1 *Ae. albopictus* had a particularly high mean F_ST_’ when compared to all other populations (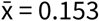; s.d. = 0.018), including substantial differentiation from Colombo-2 (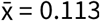 s.d. = 0.008). Colombo-2 had a much lower mean F_ST_’ when compared to all other populations (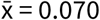; s.d. = 0.020).

**Fig 6.**
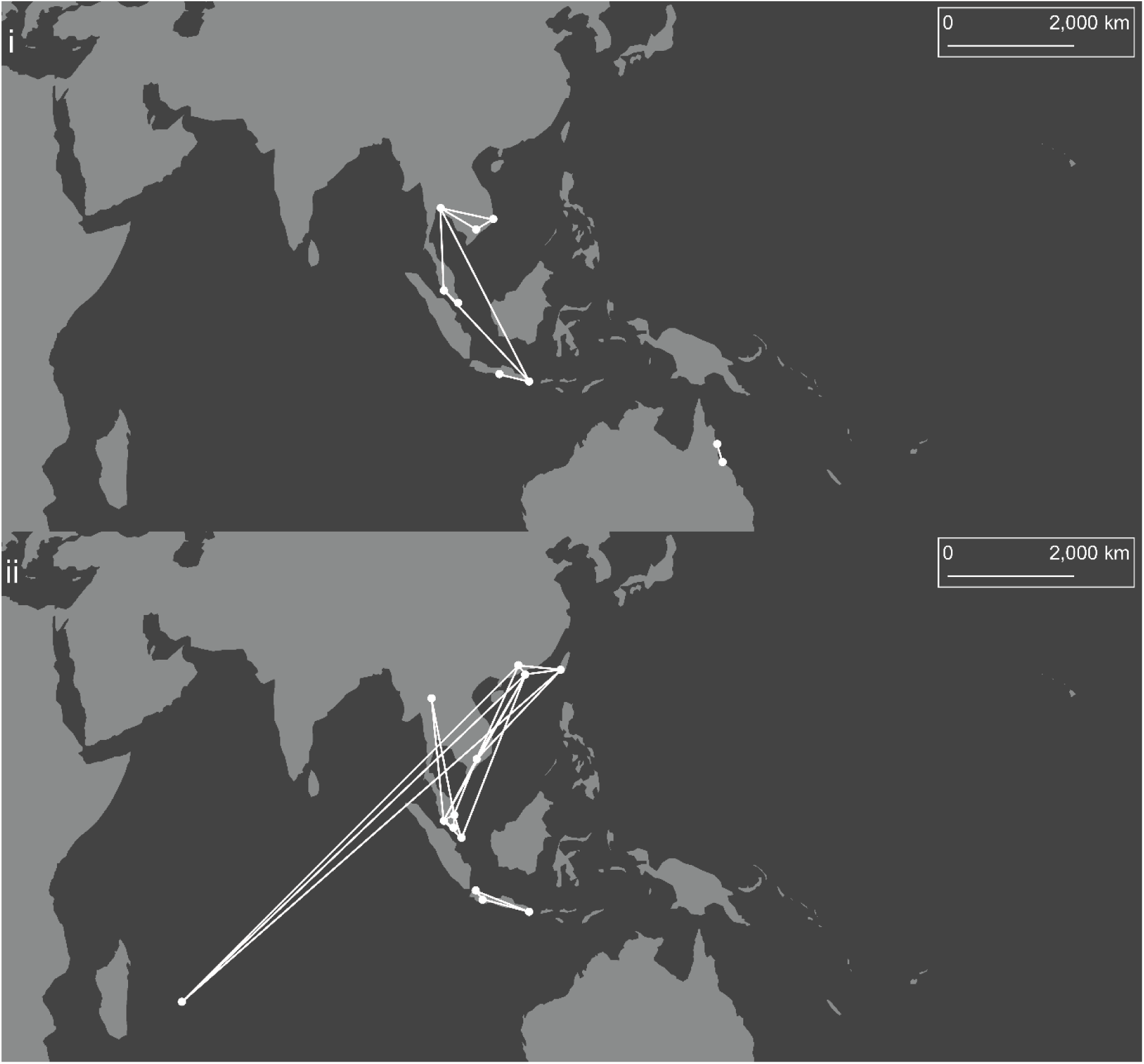
Population pairs of *Aedes aegypti* (i) and *Aedes albopictus* (ii) with mean pairwise F_ST_’ < 0.03. Circles connected by lines indicate populations with mean pairwise F_ST_’ < 0.03. The map uses a Mollweide projection with a central meridian of 120°E. The map was produced in ARCMAP 10.5.1, using shapefiles made available by www.naturalearthdata.com.

When 16 genotypes were used to calculate F_ST_’, results were similar to when 5 genotypes were used (S1 Table E). Using five genotypes tended to slightly overestimate F_ST_’, particularly for highly differentiated populations, though F_ST_’ scores were consistent relative to one another. Likewise, while using only five genotypes will lead to lower estimates of H_O_ and π than when estimated with more genotypes, H_O_ and π can be compared across populations and species. Both H_O_ and π were much higher in *Ae. aegypti* (S1 Table C) than in *Ae. albopictus* (S1 Table D).

Analysis of the folded allele frequency spectrum in *Ae. albopictus* showed that Colombo-1 had lower frequencies of rare alleles and higher frequencies of common alleles compared with other populations (Fig 7). The proportion of loci in Colombo-1 with a minor allele frequency of 0.1 was only 0.23 times the size of the mean for all populations. The proportions of loci in Colombo-1 with minor allele frequencies of 0.4 and 0.5 were 2.32 and 2.75 times higher respectively than the mean for all populations. Neither Colombo-2 nor any of the other populations consistently differed from other populations in minor allele frequency. Colombo-1 had the largest proportion of monomorphic loci out of all populations.

**Fig 7.**
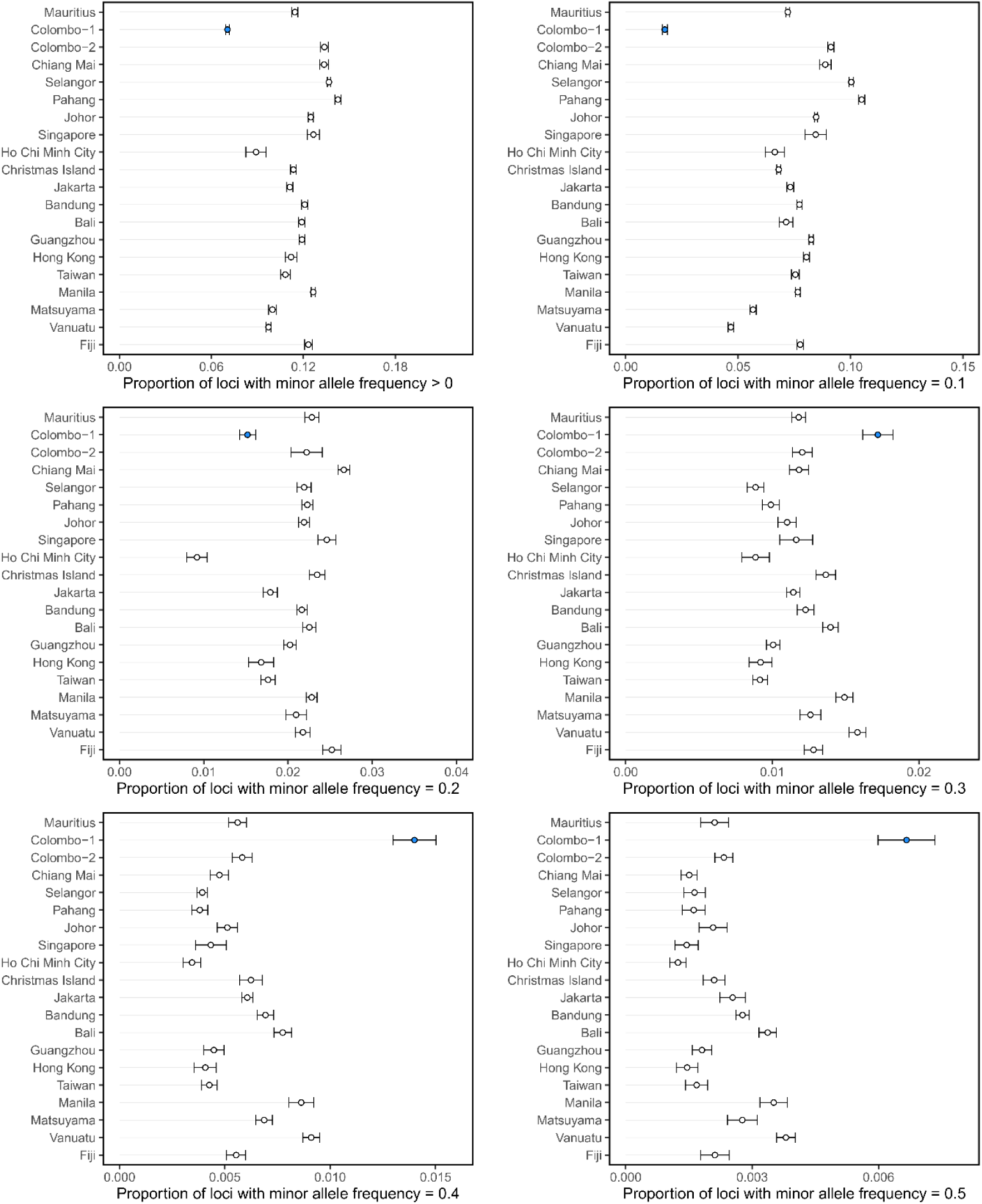
Minor allele frequencies of *Aedes albopictus* populations. Circles represent the mean frequency from the 10 subsamples, with 95% confidence intervals. The Colombo-1 population is indicated in blue.

### Landscape genomics

The dbRDAs assessing isolation by distance indicated that geographical distance was significantly associated with F_ST_’ in *Ae. aegypti* (Table 3: *F*_9_ = 2.136, *P* = 0.002) but not in *Ae. albopictus* (Table 4: *F*_9_ = 0.965, *P* = 0.534). Linear regression indicated that this relationship was positive (regression coefficient = 0.52). When each scaled vector for geographical distance was treated as an independent variable, one was significant at *P* < 0.01 in *Ae. aegypti* and none were significant in *Ae. albopictus*.

**Table 3.**
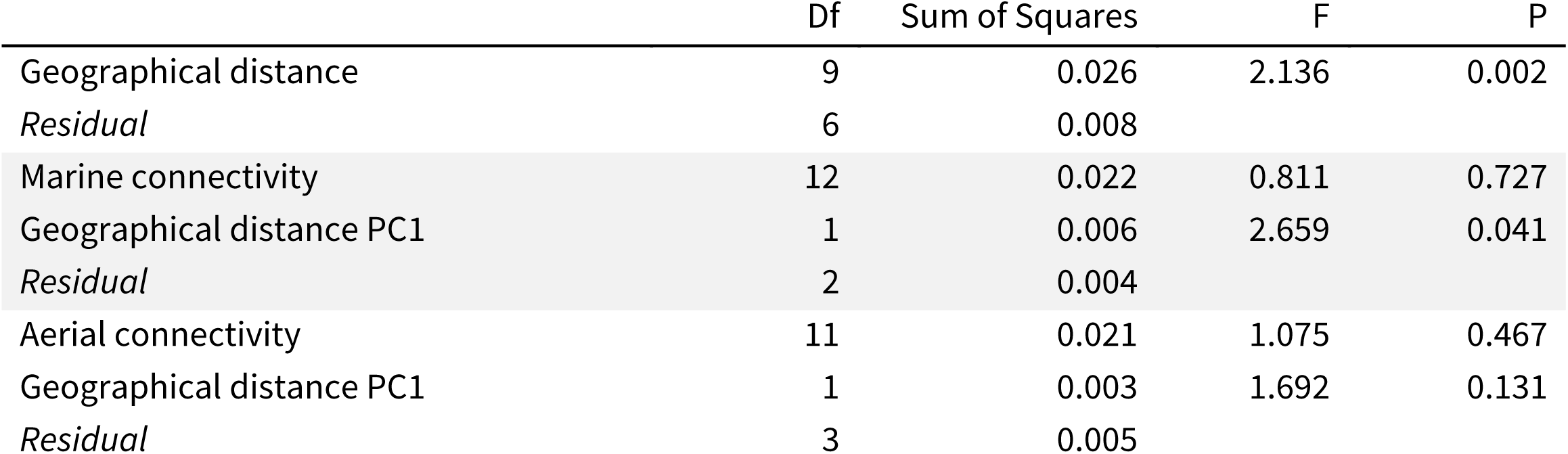
Results of dbRDAs assessing landscape genomic hypotheses in *Ae. aegypti*. All geographical distances and connectivity indices were transformed into principal coordinates (PCs) for analysis. The single PC with P < 0.01 when testing for isolation by distance was used to condition the models assessing connectivity to transportation networks.

**Table 4.** Results of dbRDAs assessing landscape genomic hypotheses in *Ae. albopictus*. All geographical distances and connectivity indices were transformed into principal coordinates (PCs) for analysis.

|  | Df | Sum of Squares | F | P |
| --- | --- | --- | --- | --- |
| Geographical distance | 9 | 0.018 | 0.965 | 0.534 |
| *Residual* | 9 | 0.019 |  |  |
| Marine connectivity | 15 | 0.034 | 2.411 | 0.034 |
| *Residual* | 3 | 0.003 |  |  |
| Aerial connectivity | 13 | 0.032 | 2.65 | 0.006 |
| *Residual* | 5 | 0.005 |  |  |

The dbRDAs assessing connectivity to transport networks indicated that F_ST_’ in *Ae. albopictus* was significantly associated with both shipping (*F*_15_ = 2.411, *P* = 0.034) and aerial (*F*_13_ = 2.650, *P* = 0.006) transportation routes (Table 4) with negative associations being detected (regression coefficients: shipping = -0.20; aerial = -0.24). Neither transport network was significantly associated with F_ST_’ in *Ae. aegypti* (Table 3), irrespective of whether the significant geographical distance vector was used to condition the models.

## Discussion

Here we present the first direct comparison of genome-wide genetic structure in the globally-invasive, dengue-vectoring mosquitoes *Ae. aegypti* and *Ae. albopictus*. Our study region, the Indo-Pacific, is the site of 70% of global dengue transmission [1], and this study contributes important knowledge to the understanding of invasion histories and ongoing gene flow within this region. While both species are already widespread in the Indo-Pacific [22], increased urbanisation and climate change may allow them to invade new regions in the coming decades [23]. Furthermore, the same dispersal processes facilitating invasions into new regions can also lead to genetic invasions of advantageous alleles into established populations, as observed recently in this region [24]. Our comparative approach indicated different genetic structure patterns in these species at each level of analysis: individual, population, and landscape genomic. For the most part, *Ae. aegypti* populations were genetically differentiated from each other, and differentiation increased with geographical distance. *Aedes albopictus* populations in the native range were split between three main regions of genetic similarity, one in East Asia, one in Southeast Asia (excluding Indonesia), and one in Indonesia. Certain populations outside the native range showed clear signs of recent invasion from these regions; specifically, Mauritius from East Asia, and Fiji and Christmas Island from Southeast Asia. *Aedes albopictus* populations in Indonesia were highly differentiated from all others, despite their proximity to other Southeast Asian populations. In Colombo, we observed two distinct *Ae. albopictus* subpopulations, one of which may represent an ancestral native-range population or cryptic subspecies. Overall these results indicate that *Aedes* spp. have established invasions in the Indo-Pacific along different pathways, and that recent gene flow patterns are different between the two species. Our findings also point to various regions of interest in the *Ae. albopictus* native range that require further investigation.

The patterns of divergence observed in *Ae. aegypti* fit with the hypothesis that *Ae. aegypti* began its invasion of the Indian Ocean region from regions to the west [5,6]. Putatively, the Indian Ocean regions and Australia were invaded from the Mediterranean, though there is some uncertainty over the timing and direction of this invasion [69,70], and the strong genetic differentiation we observed in Australian populations confounds this somewhat. Interestingly, the most highly differentiated *Ae. aegypti* populations were in the western Pacific populations of Australia, the Pacific Islands and Taiwan, suggesting that these regions do not share an invasion history with Indian Ocean regions. Separate invasions of Australia and the Pacific Islands also seem likely, based on the strong differentiation between Australian *Ae. aegypti* and those of nearby Pacific Islands such as Vanuatu and New Caledonia.

The high differentiation of western Pacific *Ae. aegypti* populations may reflect an earlier invasion of these regions, possibly from the Americas. However, these divergences could instead be due to a lack of gene flow into these populations following invasion, with strong genetic differentiation produced by the effects of drift following founder events [37]. Different structural patterns in these populations may be reinforced by high local population densities, reflecting a “founder takes all” [38] system following colonisation. Considering the strong isolation by distance observed in this species (Table 3), the relative remoteness of the Pacific Islands suggests that populations from these islands experience limited gene flow, leading to pronounced differences among them. By contrast, transport connectivity was an important influence on *Ae. albopictus* genetic structure (Table 4), and there were signs of recent dispersal into Fiji from Southeast Asia (likely Malaysia); a non-Indonesian Southeast Asian population is thus a likely source of the Fijian invasion of *Ae. albopictus* in 1989 [71]. These results require cautious interpretation, however, as most *Ae. albopictus* invasions of the Pacific are more recent than those of *Ae. aegypti*, and thus low differentiation among populations may reflect recent invasion rather than ongoing gene flow. Nevertheless, genetic similarities among geographically distant populations were more apparent in *Ae. albopictus* than in *Ae. aegypti*, and the isolation by distance patterns observed in *Ae. aegypti* indicate that gene flow in this species is more common between geographically proximate populations.

Our landscape genomics analyses point to different processes structuring *Aedes* spp. populations in the Indo-Pacific region. For *Ae. aegypti*, population structuring by geographical distance but not transportation routes accords with a pre-modern invasion timeline. *Aedes aegypti* is thought to have begun its invasion of the region in ∼18-19^th^ c. [6], a time when shipping routes likely differed from the 21^st^ century routes used in our analyses. In contrast, the more recent expansion of *Ae. albopictus* occurred at a time when international shipping routes more closely resembled the 2004 routes analysed here. Although dispersal by aircraft may be less common than by shipping in this species [25], it is worth noting that much of the material used in this study was collected near to international airports, which may exaggerate the influence of the aerial dispersal network. Also, as the metric for aerial connectivity was derived from the activity of nearby airports, aerial connectivity scores may correlate closely with other untested variables that could influence regional genetic structure, such as economic activity.

Although the observation of isolation by distance in *Ae. aegypti* but not in *Ae. albopictus* may reflect the greater impact of long-distance transportation in modern times, the capacity of *Ae. albopictus* to undergo diapause as well as quiescence [35] could be important, as mosquitoes must survive long-distance transportation for gene flow to occur. A comparison of incursion pathways of *Aedes* spp. species into Australia indicated that not only was each species dispersing from a different set of locations, but also that each species was likely transported through different vessel types, with *Ae. albopictus* found more frequently at seaports and *Ae. aegypti* at airports [25]. The most common source of *Ae. albopictus* incursions was southern China, where diapause has been observed in field populations [72]. Many other factors may also influence long-distance dispersal outcomes, such as differences in behaviour when boarding and travelling on vessels and the relative abundance of each species around airports and seaports. Better survivorship on long ship journeys may allow *Ae. albopictus* to colonise distant locations without first colonising geographically intermediate ‘stepping stones’, which accords with much of the broad-scale literature for this species [44,73]. For taxa capable of long-distance dispersal, long-range colonisation becomes more probable than stepping-stone colonisation as the required number of successful colonisations is minimised [74].

The investigation of *Ae. albopictus* in its native range revealed a pair of native-range clades covering East and Southeast Asia, with both clades found in Ho Chi Minh City (Fig 3ii). Differentiation was low between these clades (Fig 6ii). A third potential native-range clade was found in Indonesia, where populations were strongly differentiated from all other clades, confirming previous results from allozyme data [75]. Indonesia could represent the southern border of the native range, and whereas the East and Southeast Asian populations have become largely homogenised through gene flow, the Indonesian populations have remained distinct from these. Alternatively, Indonesia may have been invaded earlier than other regions. A possible ancient invasion source is India, which had considerable connectivity with Java and Bali in antiquity [76].

The *Ae. albopictus* of Colombo, Sri Lanka, appeared to represent two sympatric subpopulations: Colombo-1, a highly-divergent, ancestral subpopulation of *Ae. albopictus*, possibly a cryptic subspecies; and Colombo-2, a subpopulation seemingly produced by admixture between the ancestral Colombo-1 lineage and the Southeast Asian native-range lineage (Fig 3ii). The folded allele frequency spectrum of Colombo-1 was unlike other populations, having greater proportions of high-frequency minor alleles and fewer rare alleles (Fig 7). Several potential cryptic species closely related to *Ae. albopictus* have been identified in its Chinese native range [77], and the western extent of its range in India is not well-studied genetically. It is possible that Colombo is also within the native range of *Ae. albopictus*, and that Colombo-1 is an ancestral, native-range subpopulation. Alternatively, Colombo may have been invaded by an ancestral, native-range population from India or elsewhere. The putatively admixed Colombo-2 subpopulation is evidence of recent incursions into Colombo from the native range.

The different patterns of population genetic structure in the *Aedes* spp. described here has implications for biosecurity and biocontrol. Recently, genomics has been used to identify source locations of *Aedes* incursions into Australia and New Zealand [20,25]; these incursions threaten not only the arrival of species themselves but also the introduction of insecticide resistance alleles [24]. The genomic patterns identified here can help in identifying biosecurity threats in both species, both by revealing likely pathways of gene flow and demonstrating that long-distance marine incursions may be more likely in *Ae. albopictus* than in *Ae. aegypti*. The identification of the highly-differentiated *Ae. albopictus* clusters in Indonesia and Sri Lanka may also indicate regions where vector competence is higher or lower. For instance, *Ae. albopictus* from three populations putatively invaded from East Asia were highly susceptible to DENV-2 [78], while *Ae. albopictus* from the Torres Strait Islands, putatively invaded from Indonesia [79], had low susceptibility [80]. Genetically differentiated populations may also have differential *Wolbachia* infection status, as found in cryptic *Ae. albopictus* subspecies in China [77]. Variation in *Wolbachia* infection status may be exploitable by future dengue control efforts that involve the release of this bacterium into mosquito populations [81]. Likewise, patterns of population genetic structure can indicate areas where mosquito genetic backgrounds are likely to be similar or different to those of target populations – an important consideration for widespread species such as *Aedes* spp. that can be locally adapted to different conditions [49].

## Acknowledgements

We thank Tim Hurst, Samia Elfekih, Dayanath Meegoda, Hadian Sasmita, Nazni Wa, Pattamaporn Kittayapong, Lia Faridah, Kozo Watanabe, Thaddeus Carvajal, Chi Yung Jim, Gwendolin Wong, Ashley Callahan, Jason Axford, Stephen Doggett, Elizabeth Valerie, Craig Williams, Joe Davis and other researchers for samples. We also thank Moshe Jasper, Anthony van Rooyen, Nancy Endersby-Harshman and Nick Bell for sample processing. This research was facilitated by use of the Nectar Research Cloud, a collaborative Australian research platform supported by the Australian Research Data Commons (ARDC) and National Collaborative Research Infrastructure Strategy (NCRIS). We thank the Department of Agriculture and Water Resources (DAWR), Australian Government, for providing funding for this work. AAH was supported by Program and Fellowship grants from the National Health and Medical Research Council (NHMRC), no. 1037003. TLS and AAH were also supported by the Wellcome Trust UK, no. 108508. A-CH was funded the German Science Foundation (HO 5981/1-1) and the Swiss Tropical and Public Health Institute.

## Data Availability Statement

Aligned .bam files for 224 *Ae. aegypti* and 256 *Ae. albopictus* will be made available at the Sequence Read Archive, NCBI, by time of publication.

## Supporting Information Legends

**S1 Appendix**

This appendix contains additional information on sample processing and filtering, from the DNA extraction stage onwards.

**S1 Tables A – E**

These tables show means and upper and lower confidence ranges of: (A) pairwise F_ST_’ for *Ae. aegypti* in the Indo-Pacific region; (B) pairwise F_ST_’ for *Ae. albopictus* in the Indo-Pacific region; (C) heterozygosity and nucleotide diversity for *Ae. aegypti* in the Indo-Pacific region; (D) heterozygosity and nucleotide diversity for *Ae. albopictus* in the Indo-Pacific region; and (E) F_ST_ when calculated with five genotypes compared with when calculated with 16 genotypes.

**S1 Fig A**

**DAPCs of *Aedes aegypti* (i) and *Aedes albopictus* (ii) using lowest AIC.**

Colours indicate cluster membership. Circles are sized relative to population sample size. For *Ae. aegypti, K* = 15 was used, while *K* = 11 was used for *Ae. albopictus*. The map uses a Mollweide projection with a central meridian of 120°E. The map was produced in ARCMAP 10.5.1, using shapefiles made available by www.naturalearthdata.com.

## Author contributions

Conceptualization: TLS, AAH

Data Curation: TLS, JC

Formal Analysis: TLS, JC

Funding Acquisition: ARW, AAH

Investigation: TLS, JC, A-CH

Methodology: TLS, JC

Project Administration: TLS, ARW, AAH

Resources: TLS, A-CH, ARW, AAH

Software: TLS, JC

Supervision: AAH Validation: TLS, JC

Visualization: TLS, JC

Writing – Original Draft

Preparation: TLS

Writing – Review & Editing: TLS, JC, A-CH, ARW, AAH

## Notes

### Competing Interest Statement

The authors have declared no competing interest.

